# Artificial light at night reshapes diel brain transcriptomics in Dascyllus aruanus damselfish

**DOI:** 10.64898/2026.05.18.725701

**Authors:** Shachaf Ben-Ezra, Dana Sagi, Jasmin Lynn Mellijor, Saki Harii, Frederic Sinniger, Lior Appelbaum, Oren Levy

## Abstract

Artificial light at night (ALAN) disrupts natural light cycles and interferes with light-dependent biological processes. However, the effect of ALAN on cellular processes in wildlife is unclear. We examined diel brain transcriptomic alterations in the diurnal damselfish *Dascyllus aruanus* by comparing fish exposed to three consecutive nights of ALAN with control fish, sampled during both the day and night. ALAN partially disrupted circadian regulation transcription, altering diel expression of the core clock regulator *bmal1* and glucocorticoid-regulated genes. At night, ALAN triggered activation of genes indicative of neuronal activity and acute neural stress, along with suppression of restorative nocturnal processes. The following day, the transcriptomic divergence between ALAN-exposed and control fish expanded, with widespread downregulation of genes governing vascular homeostasis, coagulation, and immune function. Together, these findings indicate that ALAN reshapes brain transcriptomic programs across the entire diel cycle, identifying molecular signatures of physiological disruption in light-polluted marine environments.

## Introduction

Artificial light at night (ALAN) is rapidly reshaping natural light cycles, particularly in coastal regions that are experiencing increasing levels of anthropogenic night illumination due to urbanization, harbors, and tourism infrastructure. Major sources of artificial light pollution include city lights, building interior and exterior lighting, streetlights, industrial facilities, harbors and ports, and offshore oil rigs^1^. Sunlight and moonlight serve as fundamental environmental cues regulating behaviors such as foraging and reproduction^2^, as well as cellular-level diel rhythms across taxa^3^. Previous studies in terrestrial organisms and freshwater fish species have shown that nighttime illumination can disrupt circadian rhythms, endocrine function, hormonal regulation, and immunity^4–8^. However, the effects of ALAN on marine organisms, especially fish, are still poorly characterized.

Reef fish, like many other marine organisms, depend on natural light-dark cycles to regulate their daily activity patterns^9^. These fish are particularly susceptible to light pollution because they possess widespread extraocular photoreceptors^10,11^. Indeed, the research on the impacts of ALAN on marine fish has demonstrated behavioral and physiological consequences across various species and life stages. ALAN has been shown to influence reef fish foraging and feeding behavior^9,12–15^, reproduction^16^, metabolism^14,17^, growth, and survival^18,19^. However, current evidence for the influences of ALAN on behavior and physiology in marine fish often ignores cellular-level alterations. Understanding how ALAN affects reef fish gene expression is critical for establishing candidate molecular indicators for future research and assessing the broad potential biological consequences of ALAN.

In recent years, a few studies have investigated ALAN-induced gene expression changes in a range of taxa, including corals^3^, seagrass^20^, midges^21^, mosquitoes^22^, zebrafish^7^, common toads^23^, bats^24^ and laboratory rodents^25^. These studies reveal that exposure to ALAN profoundly disrupts cellular processes in both diurnal and nocturnal organisms, mostly by desynchronizing the rhythmic expression of core circadian clock genes. Additionally, ALAN triggers heightened cellular stress, inducing oxidative stress with a widespread suppression of immune function. Furthermore, light pollution commonly disrupts cellular metabolism, specifically altering lipid metabolism, energy production, and catabolic pathways. While some biological pathways are shared, the effects of ALAN on physiology and gene expression vary across species, life stages, sex, light intensity, exposure duration, tissues, and sampling time, leaving gaps in our understanding of how ALAN affects diverse life on Earth.

In this study, we investigate the transcriptomic effects of ALAN in the brain of the damselfish *Dascyllus aruanus* (Whitetail Dascyllus, *D. aruanus*), a common diurnal reef fish across the Indo-Pacific area. Using RNA sequencing and de novo transcriptome assembly, we compare gene expression profiles between ALAN-exposed and control fish brains sampled at both day and night. This approach enables us to identify the natural diel shifts in the transcriptome of *D. aruanus* fish and to detect differentially expressed genes (DEGs) at the brain tissue level. The results provide new insights into how ALAN disrupts gene expression related to circadian regulation, neuronal activity, metabolism, endocrine function, and immunity in the brains of wild reef fish. Consequently, this study contributes to a broader understanding of the impact of ALAN at the transcriptomic level.

## Results

### Brain transcriptomic response to ALAN in *D. aruanus*

Wild-caught fish obtained near Sesoko Island in Okinawa, Japan (Fig. 1A), were acclimated in two outdoor seawater flow-through tanks under natural light-dark cycles (LD, approximately 12:12 h). The fish were separated into two experimental conditions; the control group experienced natural nighttime, while the treatment group was exposed to dim ALAN (0.15-0.25 µmol m^−2^ s^−1^, 6-10 lux). Following three consecutive nights of ALAN exposure, brain samples were collected at two time points. Nighttime samples were taken one hour before sunrise (zeitgeber time, ZT23) for the control (N) and ALAN (N*) conditions. Daytime samples were collected the following day, one hour before sunset (ZT11), for the control (D) and ALAN (D*) conditions. Bulk RNA was extracted from single brains, and a de novo transcriptome was assembled, followed by transcriptomic analyses (Fig. 1B). Principal component analysis (PCA) of the 1,000 most variable genes revealed distinct clustering by gene expression profiles (Fig. 2A). Samples were grouped by sampling time, with a distinct separation between day and night groups regardless of treatment. Within these temporal clusters, samples were sub-clustered by treatment. Notably, although ALAN was applied exclusively during the night, this separation was more pronounced between the daytime groups (D* vs. D) than the nighttime groups (N* vs. N).

**Figure 1.**
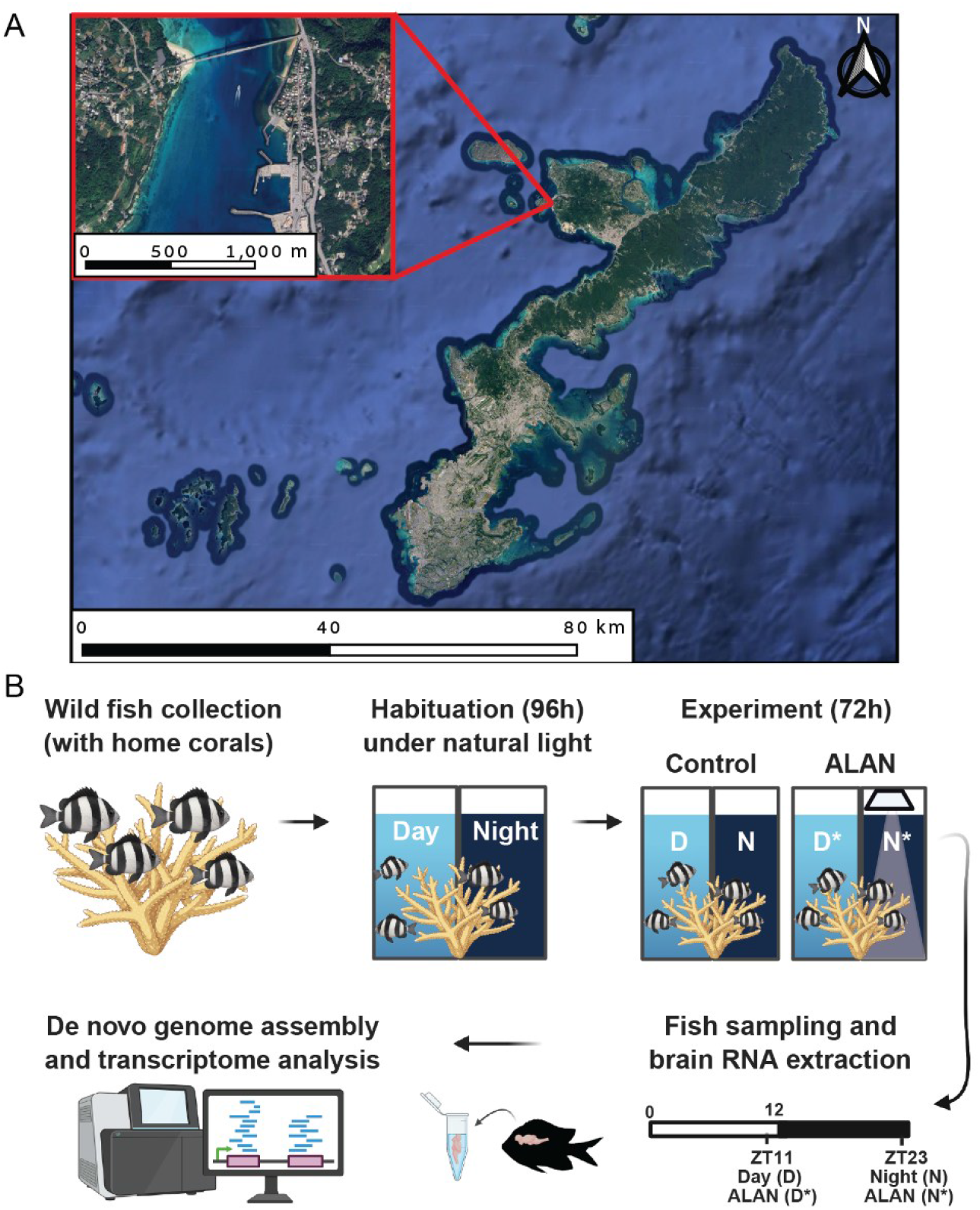
Experimental design and site location. **(a)** Collection location at the Hamasaki fishing port in Okinawa, Japan. The map was created using QGIS. Satellite imagery: ^©^ 2026 Google; Data SIO, NOAA, U.S. Navy, NGA, GEBCO; Imagery ^©^ Airbus (2015–2024). **(b)** Schematic of the experimental workflow. Wild *D. aruanus* fish were collected with their host coral colonies. Fish were habituated to laboratory conditions under a natural light-dark (LD) cycle for 96 hours. Following habituation, colonies were exposed to either a control natural LD cycle or artificial light at night (ALAN) for 72 hours. During the ALAN treatment, dim white LED light (6-10 lux) was applied from sunset to sunrise. Brains were sampled at two time points: ZT23 (one hour before sunrise) for control night (N) and ALAN night (N*) groups, and ZT11 (one hour before sunset) for control day (D) and ALAN day (D*) groups. Total RNA was extracted from brain tissue for de novo transcriptome assembly and differential expression analysis.

**Figure 2.**
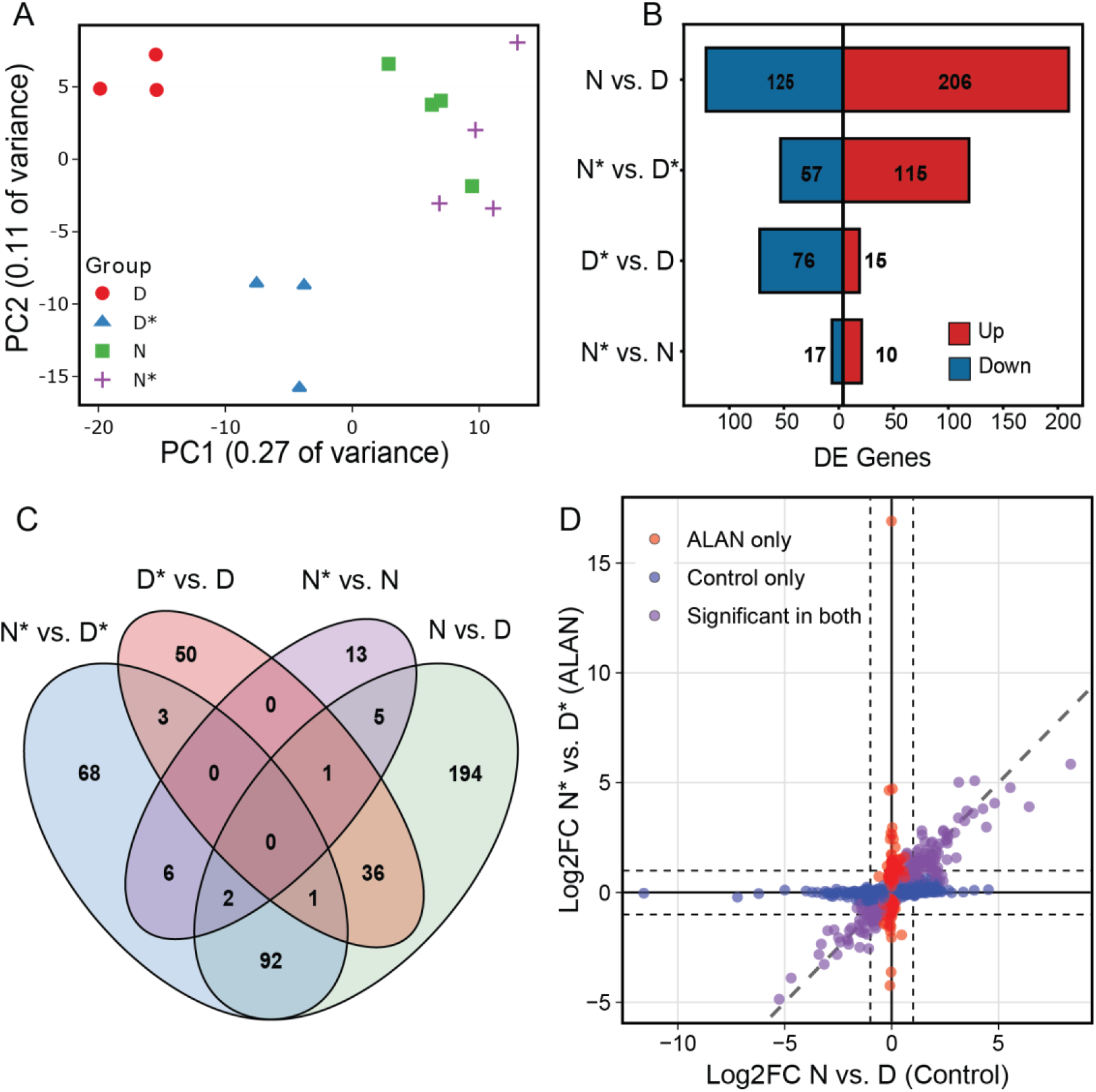
Principal component analysis (PCA) and differential expression of brain transcriptomes under control and ALAN conditions. **(a)** PCA of the 1000 most variable genes demonstrates distinct clustering by sampling time, with separation between day (D, D*) and night (N, N*) regardless of treatment. Within these temporal clusters, samples were sub-clustered by treatment. Notably, a distinct separation occurred between the daytime groups (D and D*) despite ALAN being applied exclusively at night. **(b)** Bar plot showing the number of downregulated and upregulated DEGs per pairwise comparison. **(c)** Venn diagram illustrating the overlap of DEGs among the indicated experimental comparisons. **(d)** Scatter plot of log2FC values for genes differentially expressed between day and night. Colors denote genes that are differentially expressed under ALAN only (red), control conditions only (blue), or both (purple). Vertical and horizontal dashed lines indicate the |log2FC| > 1 threshold applied in this study.

Differential expression (DE) analysis was then employed to detect the natural day-night variation (N vs. D) and to determine how ALAN modulates these diel rhythms (N* vs. D*). While natural light cycles induced substantial transcriptomic shifts comprising 331 DEGs (N vs. D), ALAN exposure masked this diel variation, resulting in only 172 DEGs (N* vs. D*). Of these, only 92 DEGs were shared with the natural control condition (Fig. 2B-D). Intriguingly, comparing natural and ALAN conditions within each time point detected a more robust transcriptomic shift during the day (D* vs. D; 91 DEGs) than during the night (N* vs. N; 27 DEGs) (Fig. 2A, B). This highlights that the cellular-level impact of nocturnal light pollution extends into the subsequent active phase.

Our results are presented in four sequential sections. First, we characterize the natural diel transcriptomic shifts under control LD conditions (N vs. D). Subsequently, we describe how ALAN alters these temporal gene expression variations (N* vs. D*). The third and fourth sections highlight genes and biological pathways that exhibit phase-specific sensitivity to ALAN at day (D* vs. D) and night (N* vs. N), respectively.

### Natural diel transcriptomics variation in *D. Aruanus* brain (N vs. D)

Natural diel differences in transcriptomic profiles were assessed by comparing fish sampled at night (N, ZT23) with those sampled during the day (D, ZT11) under control conditions, revealing 331 DEGs. Of these, 206 genes were upregulated at night, while 125 were upregulated during the day (Fig. 3A, Table S1). Gene Ontology (GO) enrichment analysis identified 104 enriched functional biological process terms (Fig. 3B, Table S2), reflecting a wide range of daily cellular transitions. As expected, the analysis identified several circadian gene groups (GO:0007623; GO:0032922; GO:0042752) (Fig. 3B) as established clock genes and clock regulators exhibited expected diel expression patterns and were among the top DEGs. Specifically, *per1, per3, ciart, cry1, nr1d1, nr1d2, sftpb, bhlhe41, bhlhe40, rpe65, ptger4*, and *cipc showed* increased expression during the night, while *clock, bmal1 (*ARNTL*) and bmal2 (*ARNTL2*), hsp90aa1, npas2, egr1*, and *nfil3* were higher during the day. Although the eyes were not sampled, transcripts associated with the visual system also showed daily phase-specific expression in brain tissue. *Arrestin-3* (*arr3*), a transcript involved in dark adaptation^26^, the retinal pigment epithelium *rpe65*^27^, and *cyp27c1*, which enhances photoreceptor sensitivity to long-wavelength light^28^, were elevated at night. Conversely, *opn3*, involved in light detection^29^, and *γ*-crystallin M2, which maintains lens integrity^30,31^, were higher during the day. These results suggest an extraretinal division of daytime and nighttime light sensing in the brain of *D. aruanus* and support the central role of photoperiodism and internal clocks in coordinating gene expression^32,33^.

**Figure 3.**
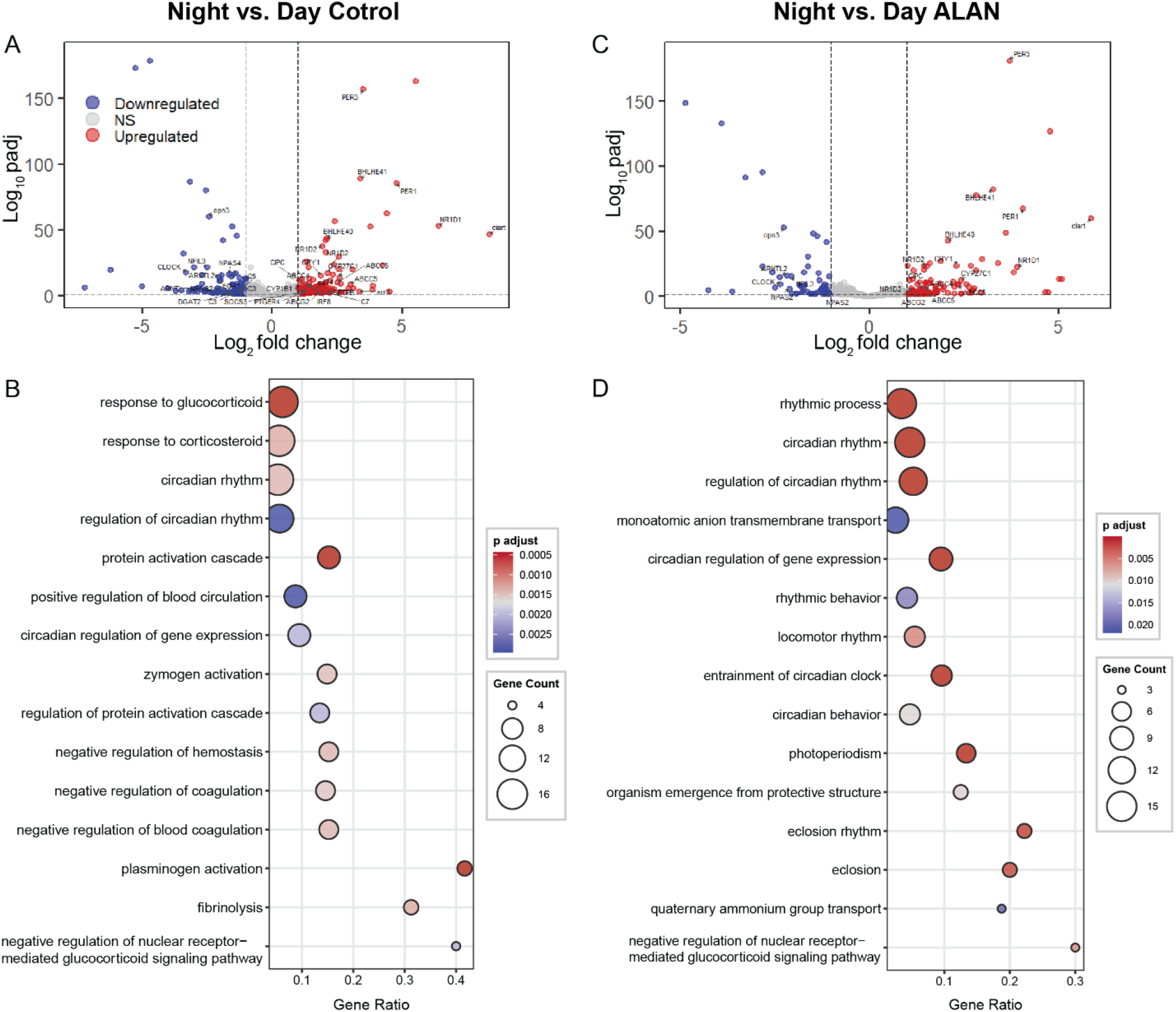
Diel expression patterns under natural and ALAN conditions. **(a, c)** Volcano plots illustrating differentially expressed genes (DEGs) between N and D (ZT23 vs. ZT11) under **(a)** control natural light and **(c)** ALAN conditions. Genes of interest discussed in the text are indicated with abbreviations. To optimize visualization, data points with log2FC > 10 were removed (resulting in the removal of one outlier in each figure). **(b**,**d)** Gene Ontology (GO) enrichment analysis of the 15 biological process terms with the lowest padj under **(b)** natural and **(d)** ALAN conditions. Terms are arranged by gene count, with the gene ratio representing the proportion of DEGs relative to the total number of genes associated with each specific GO term.

The analysis also showed enrichment of glucocorticoid-regulated genes (GO:0051384, GO:0031960) (Fig. 2B), typically via the glucocorticoid receptor (GR). ATP-binding cassette (ABC) transporters expressed in the blood-brain barrier (BBB)^34^, including *abca1, abcb1 (*also known as *mdr1*), *abcg2, abcc4*, and *abcc5*, exhibited higher expression during the night. This barrier function is complemented by the metabolic enzyme *cyp1b1*, known to be co-expressed with ABC transporters at the endothelial capillaries of the BBB^35^, which similarly displayed higher nocturnal expression. The DNA damage-inducible transcript *ddit4* (REDD1) is also regulated by the GR and was expressed more at night. *Ddit4* plays a part in the response to cellular stress, such as the accumulation of DNA damage^36^. In contrast, transcription of the neuronal PAS domain 4 (*npas4*) is suppressed by the GR^37^; accordingly, we observed that *npas4* expression was significantly higher during the day.

Other gene groups, including metabolic and immune response genes, were altered between day and night under natural conditions. Key metabolic enzymes involved in carbohydrate and energy metabolism, such as *g6pc, impa1*, and members of the malate/L-lactate dehydrogenase family, exhibited daytime upregulation. Similarly, lipid metabolism genes, including *dgat2* and *hsd17b12*, were upregulated during the day. In contrast, *prkab1*, which encodes a regulatory subunit of the AMP-activated protein kinase (AMPK), was upregulated at night. Immune markers also exhibited diel variation under natural conditions. Regulators of infection and inflammation, such as the *cd74* chaperone, the *irf8 transcription factor, ptger4* receptor, tnfsf12 and *il15 cytokines*, the *socs3* suppressor of cytokine signaling, and the complement component *c7*, were all upregulated at night. The cytokine *il15* also modulates GABA and serotonin transmission, affecting activity level and sleep^38^. Conversely, complement components *c3* and *c5* were day-upregulated. These results represent metabolic and immune-related DEGs that later lost their rhythmicity under ALAN exposure, serving as a molecular reference point for assessing how ALAN disrupts diel gene expression.

### ALAN reshapes diel variation in genes related to circadian regulation, metabolism, and immunity (N* vs. D*)

To assess the impact of ALAN on diel gene expression, we compared fish sampled at night (N*, ZT23) with those sampled the following day (D*, ZT11) under the ALAN regimen. Although these fish were still exposed to changes in light intensity between natural daylight and the dim LED light at night, ALAN exposure substantially altered the diel expression patterns observed under natural conditions. The number of genes exhibiting significant day-night differences declined from 331 (D vs. N) to 172 (D* vs. N*) (Fig. 2B-D, Fig. 3C, Table S3), indicating attenuated day-night separability and a potential rephasing or partial suppression of diel rhythms. This disruption was further evidenced by a reduction in significant GO terms, which dropped from 104 under control conditions to only 21 under ALAN (Fig. 3D, Table S4).

Core clock genes retained robust diel variation, including the oscillators *cry1, per1, per3, ciart, tef*, and clock. However, several key clock-associated regulators exhibited a marked loss or reduction in diel expression variation. The core circadian regulator *bmal1* (ARNTL), which also regulates glucocorticoid signaling, showed a reduction in diel variation, with |log2 fold change (log_2_FC)| decreasing from 1.63 in natural conditions to 0.7 under ALAN. Similarly, the immediate-early transcriptional regulator *egr1*, the circadian repressor *ciart*, and the chaperone hsp90aa1 (known to take part in *bmal1* stabilization^39,40^) all lost their diel variation under ALAN. The suppression of rhythmic signals extended to genes regulated by glucocorticoids. DEGs identified as GR-regulated under control conditions (GO:0051384), including *abcb1, cyp1b1, ddit4, npas4, socs3, agtr1, cav1, slc2a1, sftpb, ucp3, and fos*, lost their day-night variation under ALAN. Furthermore, light-sensing genes were also impacted; both *rpe65* and *arr3* failed to maintain differential expression under ALAN. Conversely, the casein kinase *csnk1d*, involved in DNA repair, apoptosis regulation, and circadian rhythms, became upregulated at night only in the presence of ALAN.

Metabolic and immune pathways were also markedly affected by nocturnal light. All metabolic and immune genes listed above (N vs. D) lost their differential expression between day and night under ALAN. Instead, a different set of genes became differentially expressed under ALAN, including genes involved in lipid signaling and synthesis, *pip5k1a, fasn*, and *ggt1*, the mitochondrial membrane respiratory chain complex I subunit *nd5*, and the alanine aminotransferase enzyme *gpt*. Notably, transmembrane transporters, including *slc10a1, slc6a13, slc22a2*, and *slc22a8 (*OAT3), widely studied drug transporters, became upregulated during the night only under ALAN. Immune response genes, such as *tagap, nlrp12*, and *igv*, also became strongly upregulated during the night under ALAN. These findings demonstrate that while core clock components may remain partially intact, downstream pathways are selectively disrupted or rephased by ALAN.

### Daytime downregulation of vascular homeostasis and immune readiness following ALAN exposure (D* vs. D)

To explore the specific cellular pathways affected by ALAN during the day, we compared transcriptomic profiles from fish sampled during the day (ZT11) following ALAN exposure (D*) with those sampled at the same time under natural control conditions (D). This comparison revealed widespread disruption of gene expression, predominantly characterized by downregulation of transcripts following exposure to ALAN. In total, 91 genes were differentially expressed, with 84% of these DEGs showing reduced expression in D* samples (Fig. 2B, Fig. S2, Table S5). Gene enrichment network analysis identified a highly connected cluster of genes associated with vascular homeostasis and immune response (Fig. 4, Table S6). This cluster was primarily driven by enrichment of genes involved in platelet degranulation (GO:0002576), negative regulation of blood coagulation and hemostasis (GO:0030195, GO:1900047), regulation of complement activation (GO:0030449), and humoral immune response (GO:0006959) (Fig. 4). These findings suggest that ALAN disrupts the anticipatory daytime physiological programs essential for vascular function and immunity.

**Figure 4.**
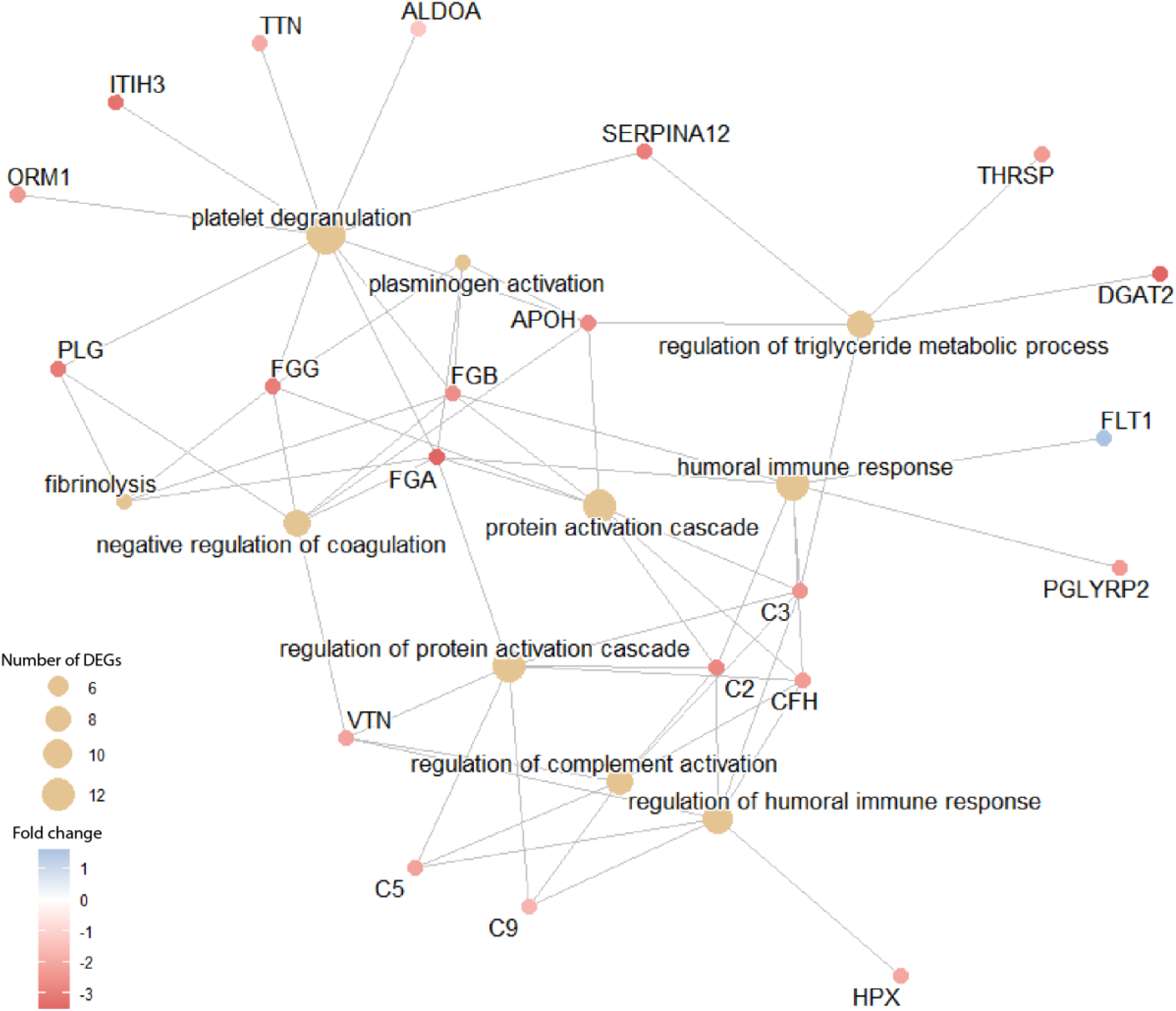
Functional enrichment network of daytime DEGs (D* vs. D). This network (cnetplot) illustrates the linkages between enriched GO terms and their associated DEGs in the daytime comparison (ZT11) between ALAN and control conditions (D* vs. D). The 10 most significant GO terms are presented, with redundant terms removed for visual clarity. Large beige nodes represent specific biological processes, where node size is proportional to the total number of genes enriched within that pathway. Smaller colored nodes represent individual genes; the color gradient reflects the observed log_2_FC, with red indicating downregulation and blue indicating upregulation under ALAN.

Only 15 genes were upregulated by exposure to ALAN at daytime (Fig. 2B). Among these few differentially expressed genes were the angiogenesis regulators *eltd1,rgs5, and flt1*, which is also involved in neuroprotective hormonal immune responses^41^. Collectively, these genes often serve as indicators and enhancers of tumor formation^42,43^. Additionally, expression of *prl*, the neuroendocrine hormone gene prolactin, was found to increase under ALAN. In teleosts, *prl* is known to be influenced by photoperiod and stress, and its surge in ALAN-exposed fish suggests a compensatory endocrine response and altered pituitary gland activity^44^. We also observed an upregulation of the gene *nr1d1*, which encodes the nuclear receptor *Rev-erbα*, indicating a phase shift or compensatory response in the circadian clock mechanism^45^.

The daytime transcriptome further revealed suppression of the physiological machinery governing blood homeostasis and vascular integrity under ALAN. Transcripts for the three fibrinogen chains, *fga, fgb*, and *fgg*, were markedly downregulated, potentially compromising the core components essential for clot formation. This inhibitory effect extended to key regulators of the coagulation cascade and fibrinolysis, including *plg, apoh*, and *vtn*. Furthermore, ALAN led to daytime downregulation of genes related to platelet degranulation, such as *serpina12, itih3, orm1, ttn, and aldoa*. Given that platelets are vital for both systemic coagulation and brain function, the concurrent suppression of these pathways suggests that nocturnal light exposure may disrupt broader neuroinflammatory and neurodegenerative processes^46^. Indeed, among the most significantly suppressed transcripts were genes involved in inflammation and cell signaling. The core inflammasome component *nlrp12* was strongly downregulated, alongside reduced transcripts for complement components c2, *c3, c5, c9*, and *cfh*. Several metabolic genes followed a similar trend, including *thrsp*, which encodes the thyroid hormone-responsive protein involved in lipid metabolism, and *hpx*, which facilitates heme and iron transport^47^. Finally, ALAN disrupted the tyrosine catabolic pathway by downregulating *fah* and *hpd*, enzymes that provide biochemical precursors for essential neurotransmitters such as dopamine and epinephrine, as well as the hormone thyroxine. Notably, *fah* is also associated with the regulation of sleep-wake patterns, further linking these metabolic shifts to broader behavioral and physiological disruption^48^.

### ALAN exposure elicits nocturnal stress and arousal in *D. aruanus* (N* vs. N)

To assess the impact of ALAN on nighttime gene expression, we compared transcriptomic profiles from fish sampled during the night under ALAN exposure (N*, ZT23) with those sampled at the same time under control conditions (N). Although this comparison yielded the fewest DEGs, the observed 27 DEGs provide critical information regarding the immediate neural response to nocturnal light (Fig. 5, Table S7). GO enrichment analysis identified a single significant biological process term: response to mineralocorticoid (GO:0051385), enriched by the induction of *npas4, fos*, and *rpl27*.

**Figure 5.**
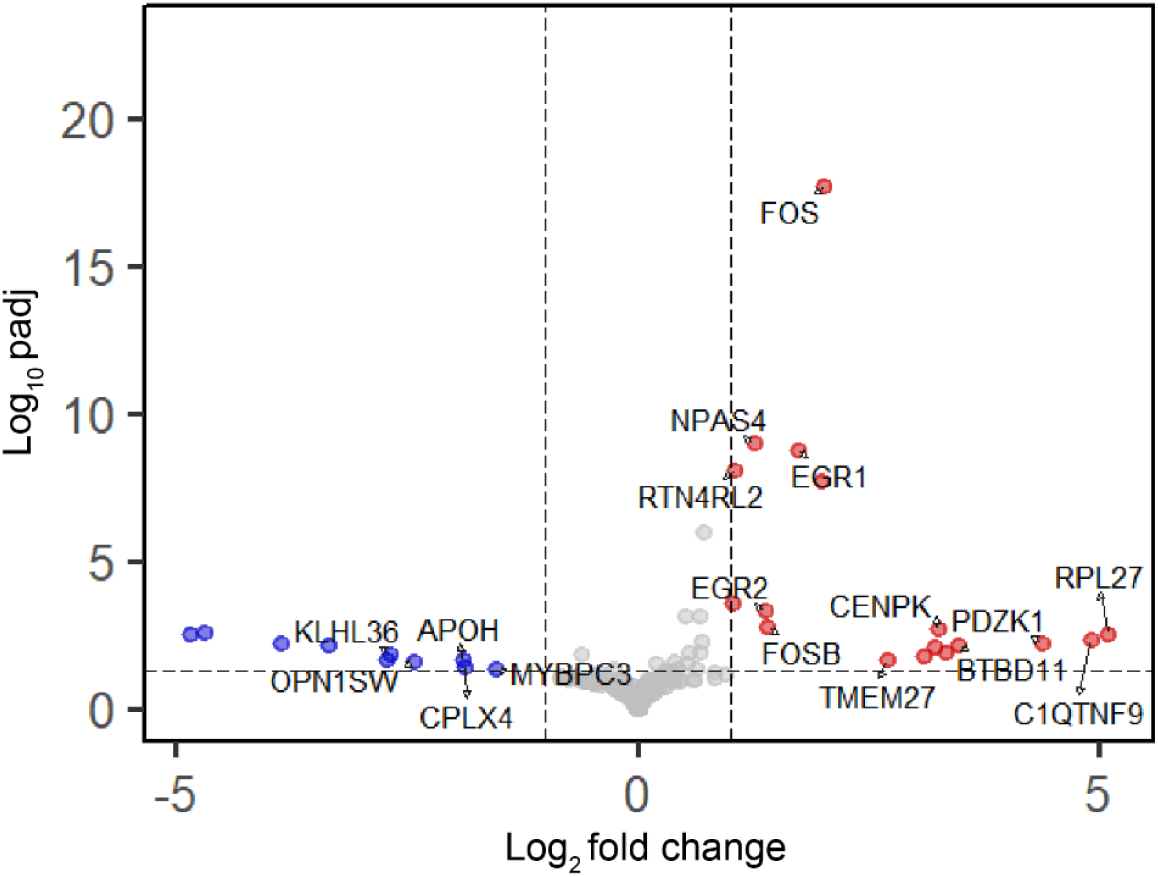
Volcano plot of nighttime DEGs (N* vs. N). Annotated volcano plot illustrating DEGs in brain samples collected at the end of the night (ZT23) under ALAN compared to the control light regimen (N* vs. N). Red and blue points represent genes with a |log2FC| > 1 and padj < 0.05, indicating upregulation and downregulation under ALAN, respectively. This comparison reveals the immediate neural response to nocturnal light, characterized by the induction of immediate-early transcription factors.

A prominent feature of ALAN exposure was the induction of immediate-early transcription factors, which serve as markers of neural activity and acute stimulation. Specifically, *fos, fosb, egr1*, and *egr2*, were strongly upregulated during the illuminated night compared to the control night. This coordinated response included upregulation of additional activity-dependent transcripts, such as *npas4*, which is involved in maintaining excitatory-inhibitory balance^37^; *btbd11*, a regulator of glutamatergic synapses and neuronal circuit function^49^; and *rtn4rl2*, involved in transsynaptic signalling and regulation of synapse development^50^. This activation pattern is characteristic of acute neural stress and heightened arousal, suggesting that ALAN triggers a state of nocturnal alertness in *D. aruanus*. Beyond activity-dependent markers, the gene *cenpk*, important for mitotic progression^51^, together with transmembrane transport regulators such as *tmem27*, and *pdzk1*, showed increased expression under ALAN. In addition, ALAN exposure induced the overexpression of *interleukin-1 receptor-like* transcript, implicating activation of inflammatory signaling. Conversely, ALAN repressed the expression of several genes essential for nocturnal restorative functions and sensory perception. These included *opn1sw (sws1)*, a short-wavelength-sensitive opsin usually involved in cone photoreception^52^, the synaptic vesicle-associated genes *cplx4*^53,54^, and *apoh*, a circadian rhythmically expressed gene important for lipoprotein metabolism, coagulation, and hemostasis^55^. Overall, these findings support the conclusion that ALAN disrupts nocturnal neurological function, reactivating daytime-like neuroactivity programs while suppressing restorative pathways.

## Discussion

ALAN disrupts the fundamental circadian rhythms and chronophysiology of organisms. In this study, we demonstrate that ALAN induces a profound reorganization of brain gene expression in the diurnal reef fish *D. aruanus*, altering both nocturnal and diurnal transcriptomic states. Under natural light conditions, the brain transcriptome exhibits a coordinated division of cellular activity between day and night. However, ALAN exposure dampens this intrinsic regulation at night and extends transcriptional disruption into the subsequent active phase. To consolidate these effects, we performed comparative KEGG enrichment analyses across all comparisons, revealing coherent functional shifts (Fig. 6). These findings indicate that light pollution compromises essential pathways, ranging from circadian regulation and neuronal activity to brain vascular homeostasis and immune readiness.

**Figure 6.**
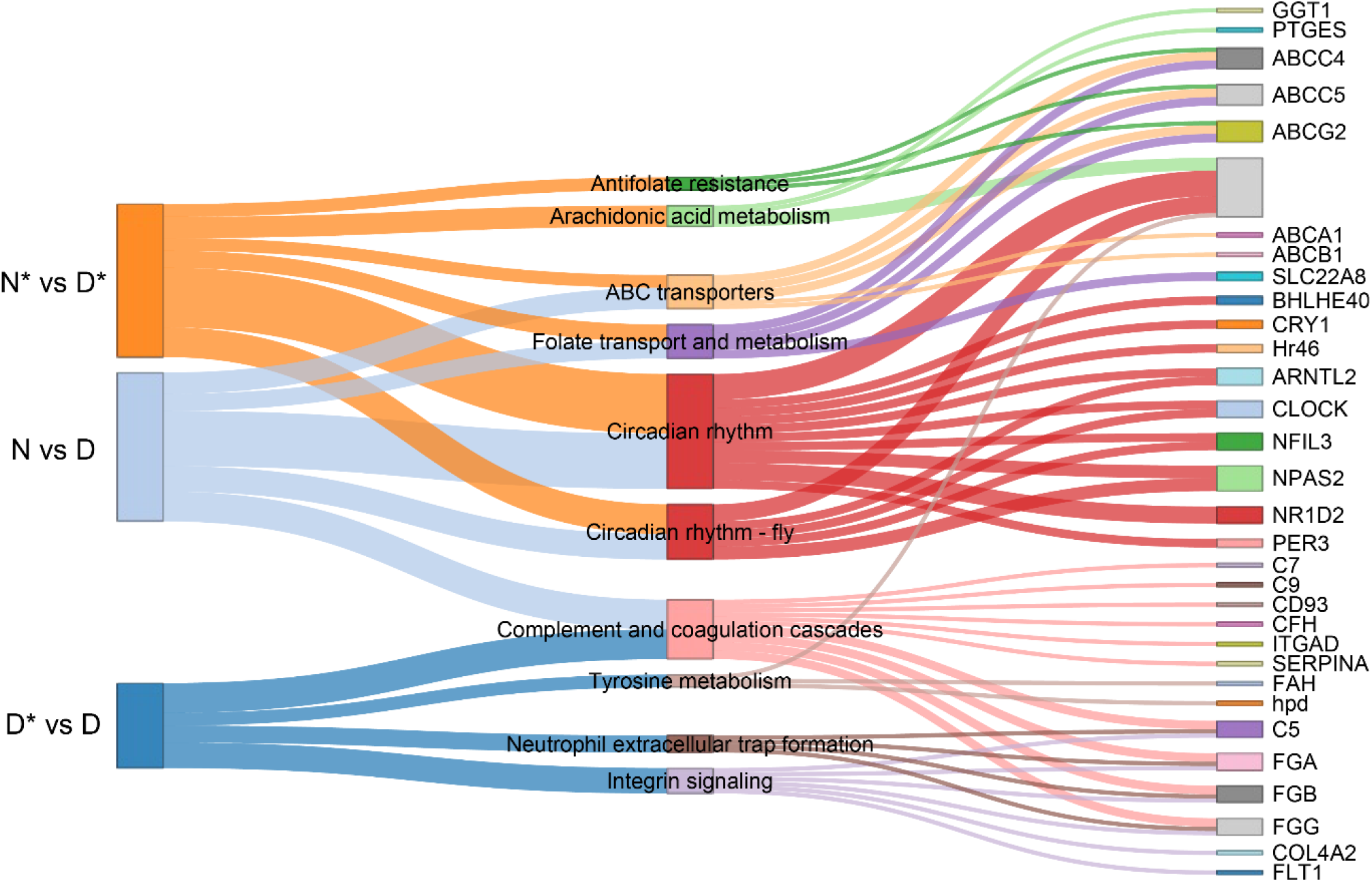
Integrated gene-to-pathway associations across diel and ALAN comparisons. Alluvial plot illustrating the results of KEGG enrichment analysis and differentially expressed genes (DEGs) across three experimental comparisons: natural night vs. day (N vs. D), ALAN night vs. day (N* vs. D*), and ALAN day vs. natural day (D* vs. D). Note that the comparison between ALAN night and natural night (N* vs. N) did not yield significantly enriched KEGG functional pathways. Left nodes represent the three differential expression comparisons. Middle nodes represent significantly enriched KEGG pathways, with node and alluvia width proportional to the number of genes contributing to that specific functional category. Right nodes identify specific constituent genes; the cumulative width of a gene node reflects its presence across multiple pathways or comparisons.

The robust day-night clustering of gene expression observed under natural light reflects a coordinated division of cellular activity governed by light sensing and internal oscillators^33^. Beyond the visual system, changes in photoreceptor gene expression support the existence of direct light sensing within the fish brain^56,57^. Light pollution has previously been shown to desynchronize the cellular expression of circadian regulators in the brains of zebrafish^7,58^, *Drosophila* and rodents^59^ following both acute and prolonged exposure. After three days of ALAN exposure, our data indicate that the core clock mechanism in the *D. aruanus* brain remained generally intact. This aligns with the characteristic of clock oscillators that maintain rhythmic expression even in the absence or masking of environmental diel cues^60^. However, an initial desynchronization of clock regulators occurred, primarily evident in *bmal1* expression and its associated regulators, *nr1d1 and hsp90aa1*. As a positive transcription factor, *Bmal1* drives the rhythmic expression of *per* and *cry* genes through a transcription-translation feedback loop^61,62^. A deficiency in *bmal1* expression carries broad implications for steroid signaling and endocrine function^63^. Indeed, ALAN chronodisruption extends to endocrine-mediated transcriptional regulation involving glucocorticoid signaling. Endocrine regulation in fish is driven by circadian and stress-induced cortisol^64^, which interacts with melatonin to preserve physiological rhythms^65^. Under natural conditions, we observed clear diel rhythms in GR-regulated genes, aligning with the anticipatory cortisol peaks typical of diurnal species^66^. However, ALAN suppressed this rhythmic expression, suggesting a loss of cortisol or GR signaling. Previous studies in fish have shown no robust changes in circulating cortisol under ALAN, but rather demonstrate a suppression of melatonin that can inhibit GR nuclear activity^4,65^. Our results highlight the need to further investigate how nocturnal light compromises the intersection of circadian clocks, glucocorticoid regulation, and GR signaling.

ALAN exposure acutely altered cellular resting and sleep signatures in the brain, affecting the expression of genes involved in immediate responses to stimuli, such as *fos, fosb*, and *egr1*^67^, as well as regulators of synaptic plasticity (*npas4* and *btbd11*)^49^ and transsynaptic signaling (*rtn4rl2* and cplx4)^50^. These changes correlate with classical molecular responses to acute neural and light stimulation^68^ and suggest that ALAN triggers a state of heightened nocturnal alertness, reactivating daytime-like neuronal transcripts and inducing neural stress. Accordingly, ALAN exposure extensively reshapes the neuroimmune landscape, inducing nighttime inflammatory signaling followed by daytime immune suppression and a loss of diel immune regulation. Daytime downregulation of key immune transcripts, including *nlrp12* and complement cascade genes (*c2, c3, c5, c9, cfh*), alongside disrupted diel rhythmic expression of immune regulators (*cd74, il15, irf8, socs3*), suggests an inflammatory response that is consistent with evidence of ALAN-induced neuroinflammation in vertebrates^69,70^. Together, these patterns indicate that ALAN promotes a non-restorative resting phase and a stress-immune response that may increase susceptibility to pathogens during the active period^71^. Additionally, these transcriptomic alterations, including circadian clock dysregulation, neuroinflammation, and neural function, closely parallel the known transcriptomic alterations of reef fish under ocean acidification^72^. Therefore, nocturnal light pollution may compromise a fish’s capacity to tolerate and adapt to concurrent environmental pressures.

ALAN daytime transcriptomic disruptions under ALAN have major implications for brain vascular integrity and blood homeostasis. The observed diel variation in BBB efflux transporters under natural light conditions supports the critical role of circadian rhythms in regulating the BBB^73^. ALAN altered the expression of genes maintaining BBB integrity, including the active efflux transporter *abcb1* and the *cyp1b1* enzyme. Additionally, daytime suppression of fibrinogen chains, clot breakdown modulators, and platelet degranulation factors indicates a potential compromise in systemic coagulation. Given the vital role of platelets in brain function, this suppression may correlate with broader neuroinflammatory and neurodegenerative disruption^46^. Daytime vascular disruptions are further evidenced by upregulation of angiogenesis regulators, which can indicate tissue stress, immune distraction, or, in some cases, tumor formation^41–43^. These transcriptomic shifts align with recent findings in mouse models, where short-term ALAN altered hippocampal vascular networks and connectivity, following alterations in transcripts associated with angiogenesis and blood-brain barrier maintenance^25^. These findings collectively suggest that ALAN disrupts the genetic landscape required for vascular homeostasis and neurodegenerative processes.

Taken together, our results suggest that ALAN imposes a sustained allostatic load, which disrupts the biological night and reprograms daytime gene expression. These findings add to the growing evidence that ALAN drives widespread physiological dysfunction across taxa and provide a foundation for necessary follow-up research. Studies incorporating multiple sampling points across the diel cycle are required to reveal whether the observed temporal shifts stem from amplitude loss or phase shifts of rhythmic genes^7^. Additionally, dedicated studies should further establish whether these shifts scale with ALAN exposure intensity, spectrum, and duration. Importantly, it is also necessary to determine how they translate into compromised fitness in wild populations. The genetic signatures reported here establish candidate molecular indicators to motivate follow-up cellular, endocrine, and behavioral assays. Such work will determine the full scale of light pollution implications for humans and wildlife.

## Methods

### Fish collection and experimental design

To investigate the transcriptional response to artificial light at night (ALAN) in the reef fish *Dascyllus aruanus*, we conducted a controlled laboratory experiment in Okinawa, Japan during March 2023. Six colonies of *D. aruanus* and their host coral colonies were collected from the Hamasaki fishing port in Okinawa. Corals were collected under Okinawa Prefecture permit No. 4-28. All coral colonies were returned to their original collection site after the experiment. The colonies were maintained in two separate seawater flow-through tanks (three colonies per tank), under natural daylight, with a shading net above the tanks, at the Sesoko Marine Research Station of the University of the Ryukyus in Sesoko Island, Okinawa, Japan. Following collection, the fish were allowed to recover and acclimate for 96 hours under natural light-dark (LD) conditions (day length of 11 hours and 45 minutes). After the habituation phase, experimental light treatments were applied for 72 hours. One tank served as a control group and remained under natural LD conditions during a full moon. The second tank was subjected to ALAN using white LED light (6500 K; spectral range 400-700 nm), activated daily at sunset and turned off at sunrise. The irradiance at the water surface during ALAN exposure was maintained at 0.15-0.25 µmol m^−2^ s^−1^, measured using a LI-193 underwater Spherical Quantum Sensor and Ocean Optics EVERFINE SPIC-200BW. This light spectrum and intensity were chosen to match the most common lighting used in modern coastal and urban areas. These parameters were based on light intensities measured at coral reefs that lie in proximity to urbanized coastal areas to ensure high ecological relevance^74,75^.

### Sampling fish brains

Fish sampling occurred after three consecutive nights of treatment. We sampled at night (ZT23, one hour before sunrise) for control (N) and ALAN-exposed night (N*) conditions, and on the subsequent day (ZT11, one hour before sunset) for control (D) and ALAN-exposed day (D*) conditions. All sampled fish were immediately euthanized using clove oil and placed on ice while their size was measured. Fish were decapitated, and brains were preserved in RNAprotect Tissue Reagent (Qiagen, Hilden, Germany; Cat. No. 76104) until RNA extraction.

### RNA extraction

Total RNA was extracted from individual brain tissues of *D. aruanus* using the Direct-zol™ RNA Miniprep kit (Zymo Research, USA), following the manufacturer’s protocol. Briefly, each brain sample was homogenized in 200 µL of TRIzol™ reagent (Invitrogen, USA) using an electric pestle homogenizer. Lysates were cleared by centrifugation at 16,000 × g for 30 seconds at 4 °C. The supernatant was mixed with ethanol and processed according to the Direct-zol™ protocol, including on-column DNase I treatment. RNA was eluted in 50 µL of RNase-free water. RNA concentration and quality were assessed using a Qubit Fluorometer and an Agilent 2200 TapeStation (RIN>7).

### De novo transcriptome assembly and annotation

Three RNA samples representing different experimental conditions (D, N, N*) were used for de novo transcriptome assembly. Libraries were prepared using the TruSeq Stranded mRNA Library Prep Kit (Illumina) and sequenced on a NovaSeq 6000 system (Illumina) with an SP 300-cycle paired-end run. After demultiplexing, adapter and quality trimming were performed with Trimmomatic v0.39^76^, yielding 92M, 93M, and 101M read pairs for D, N, and N*, respectively. Two de novo assemblies were generated using Trinity v2^77^: one from all three samples and one from N* only. The --SS_lib_type parameter was set to RF to retain strand-specific information, improving the identification of overlapping ORFs. Assembly quality was assessed via BUSCO v5^78^ and by mapping trimmed reads back to the respective assemblies. Potential contamination in the combined assembly was evaluated using BLASTn (NCBI BLAST+)^79^ to assign taxonomic identity to contigs. The vast majority of contigs had Stegastes partitus as their first BLAST match (Fig. S1). Stegastes partitus is the most closely related species with a full genome available in NCBI. To generate a non-redundant transcript set, super-transcripts were constructed using Trinity_gene_splice_modeler, selecting one representative isoform per gene.

Following de novo assembly, 14 RNA samples were processed using the INCPM mRNA-seq library preparation kit (non-stranded). Libraries were multiplexed with the stranded samples and sequenced in the same NovaSeq 6000 SP flow cell (Illumina), generating between 26 and 40 million paired-end reads per sample. Specifically, sequencing depth ranged from 26M (ED2) to 40M (MD6) reads per sample, with most samples yielding between 31M and 36M reads. Approximately 9-11 million read pairs per sample were successfully mapped to the 40,000 longest annotated contigs in the reference transcriptome. Adapter and quality trimming were conducted using Trimmomatic v0.39, applying the same parameters as for the stranded samples. Reads were aligned to the reference transcriptome and quantified using RSEM^80^, as implemented in the Trinity utility abundance_estimates_to_matrix, with Bowtie2 ^81^used for alignment. Functional annotation was performed with eggNOG-mapper v2.1.7^82^, using the eggNOG v5 database^83^ in CDS mode and employing Diamond^84^ for sequence alignment. For downstream analyses, only the longest 40,000 contigs with eggNOG annotation from the combined assembly were retained. Sequencing and primary data processing were conducted at the Mantoux Bioinformatics Institute of the Nancy and Stephen Grand Israel National Center for Personalized Medicine, Weizmann Institute of Science (Rehovot, Israel).

### Differential expression, gene ontology (GO), and Kyoto Encyclopedia of Genes and Genomes (KEGG) enrichment analysis

Differential gene expression analysis was performed using the DESeq2 package in R^85^. Raw read counts were used as input, and DESeq2 performed internal normalization by estimating size factors using the median-of-ratios method. DEGs were tested using the negative binomial generalized linear model (GLM) and the Wald test. To account for high dispersion in low-count genes, log2FC estimates were refined using the adaptive shrinkage estimator with the R package ashr^86^. P-values were adjusted for multiple testing using the Benjamini-Hochberg procedure to control the false discovery rate (FDR). Genes were defined as differentially expressed (DE) based on an adjusted p-value (padj) < 0.05 and an |log2FC| > 1. To filter out low-abundance transcripts per comparison, we required a maximum raw count of ≥30 reads and ≥10 reads in at least two samples. GO enrichment analysis was performed using the R package clusterProfiler (v4.18.4)^87^ and visualized using enrichplot (v1.32.0)^88^. The analysis was restricted to the “Biological Process” ontology with terms containing at least 3 significant genes. Terms with a Benjamini-Hochberg padj < 0.05 were considered significant. Network visualizations (cnetplot) were generated using clusterProfiler to illustrate the linkage between specific genes and their associated biological processes. Similar to GO term analysis, KEGG pathway enrichment was performed on differentially expressed genes, where pathways with padj < 0.05 were considered significant. Significantly enriched pathways were then visualized using network3D^89^.

## Resource availability

RNA-seq raw data and the de novo transcriptome assemblies have been deposited in NCBI, GEO repository number GSE331185.

## Acknowledgements

We thank the staff and students at the Sesoko Marine Research Station, University of the Ryukyus, for their hospitality and support during fieldwork. We are particularly grateful to the members of the laboratories led by Prof. Saki Harii and to Prof. Ezri Tarazi for their essential field assistance in Okinawa. Furthermore, we thank Dr. Amir Szitenberg and Dr. Eviatar Weizman from the Mantoux Bioinformatics Institute at the Weizmann Institute of Science, and Dr. Tirza Doniger at the Faculty of Life Sciences, Bar Ilan University, for their valuable technical and computational assistance with transcriptome analysis. We thank Dr. Sarit Lampert and Dr. Orly Yaron at the Kanbar Facility for Core Research Equipment, Bar Ilan University, for their guidance regarding RNA extraction. Finally, we thank the members of the Levy and Appelbaum laboratories for their assistance and support with the molecular and computational work. This study partially fulfills the requirements for the PhD thesis of S.B.E. at the Faculty of Life Sciences, Bar Ilan University, Israel.

## Author contributions

Conceptualization-SBE, DS, OL, LA; Data curation-SBE, DS; Formal Analysis-SBE, DS, JLM; Funding acquisition-LA, OL, SH, SBE; Investigation-SBE, SH, FS, OL; Methodology-SBE, DS, SH, FS, LA, OL; Project administration-SBE, OL, LA, SH, FS; Resources-SH, FS, OL, LA; Software-SBE, DS, JLM; Supervision-OL, LA; Validation-SBE, LA, OL; Visualization-SBE, DS, JLM; Writing - original draft-SBE, DS, OL, LA; Writing - review & editing-SBE, DS, OL, LA, SH, FS

