## Supplementary Materials for "Artificial light at night reshapes diel brain transcriptomics in Dascyllus aruanus damselfish"

**List of supplementary materials:**

Figures S1 to S2
Tables S1-S7

**Supplementary figures:**

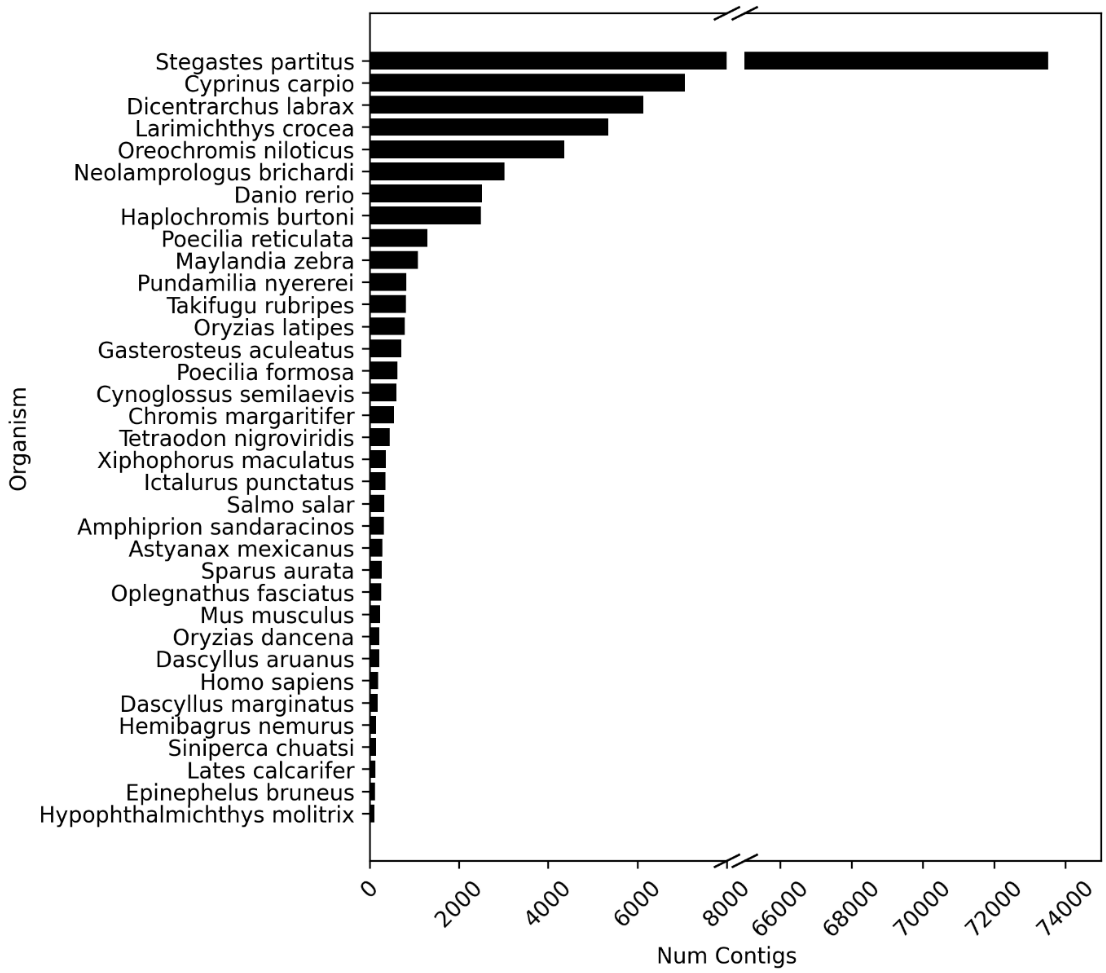


**Figure S1. Taxonomic assignment of assembled contigs.** Bar plot representing the number of contigs assigned to various organisms following BLASTn (NCBI BLAST+) analysis. The vast majority of contigs matched to *Stegastes partitus* as their primary BLAST hit. *Stegastes partitus* is the most closely related species with a fully sequenced genome available in the NCBI database.

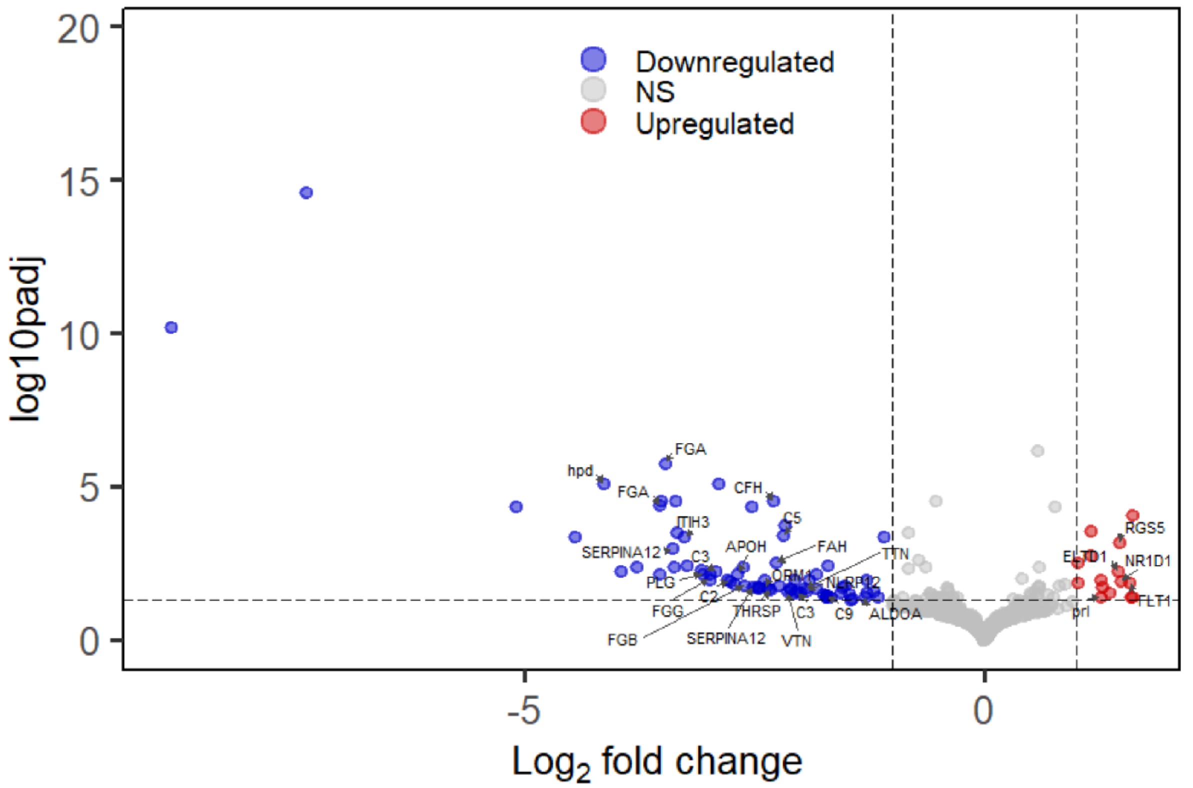


**Figure S2. Transcriptomic shifts in daytime brain samples following nocturnal ALAN exposure.** Annotated volcano plot illustrating differentially expressed genes (DEGs) in daytime samples (ZT11) under ALAN compared to the control light regimen (D* vs. D). Points are colored to indicate genes that are significantly downregulated (blue) or upregulated (red) under ALAN. The plot highlights a widespread suppression of transcripts. To optimize visualization, data points with log2FC > 10 were removed (resulting in the removal of one outlier).

**Supplementary tables (included as one file with separate sheets):**

**Table S1:** Control treatment day vs. night DEGs (D vs. N)

**Table S2:** Control treatment day vs. night GO enrichment results (D vs. N)

**Table S3:** ALAN treatment day vs. night DEGs (D* vs. N*)

**Table S4:** ALAN treatment day vs. night GO enrichment results (D* vs. N*)

**Table S5:** Daytime DEGs (D* vs D)

**Table S6:** Daytime GO enrichment results (D* vs D)

**Table S7:** Nighttime DEGs (N* vs. N)
